# The potential role of miR-21-3p in coronavirus-host interplay

**DOI:** 10.1101/2020.07.03.184846

**Authors:** Stepan Nersisyan, Narek Engibaryan, Aleksandra Gorbonos, Ksenia Kirdey, Alexey Makhonin, Alexander Tonevitsky

**Author notes:** Corresponding author: Stepan Nersisyan.

## Abstract

Host miRNAs are known as important regulators of virus replication and pathogenesis. They can interact with various viruses by several possible mechanisms including direct binding the viral RNA. Identification of human miRNAs involved in coronavirus-host interplay is becoming important due to the ongoing COVID-19 pandemic. In this work we performed computational prediction of high-confidence direct interactions between miRNAs and seven human coronavirus RNAs. In order to uncover the entire miRNA-virus interplay we further analyzed lungs miRNome of SARS-CoV infected mice using publicly available miRNA sequencing data. We found that miRNA miR-21-3p has the largest probability of binding the human coronavirus RNAs and being dramatically up-regulated in mouse lungs during infection induced by SARS-CoV. Further bioinformatic analysis of binding sites revealed high conservativity of miR-21-3p binding regions within RNAs of human coronaviruses and their strains.

## INTRODUCTION

Coronavirus disease 2019 (COVID-19) caused by the severe acute respiratory syndrome coronavirus 2 (SARS-CoV-2) acquired pandemic status on March 11, 2020 making a dramatic impact on the health of millions of people (Zhou et al., 2020a; Remuzzi and Remuzzi, 2020). Lung failure induced by the acute respiratory distress syndrome (ARDS) is the most common cause of death during viral infection (Xu et al., 2020).

MicroRNAs (miRNAs) are short (22 nucleotides in average) non-coding RNAs which appear to regulate at least one-third of all human-protein coding genes (Nilsen, 2007). Namely, assembling with a set of proteins miRNA forms an RNA-induced silencing complex (RISC) and binds the 3’-UTR of the target mRNA. The latter promotes translation repression or even mRNA degradation (Carthew and Sontheimer, 2009). Multiple works suggest the critical function of miRNAs in the pathogenesis of various human diseases. Thus, alteration of miRNAs expression is observed during different types of cancer (Di Leva et al., 2014; Shkurnikov et al., 2019), cardiovascular (Schulte et al., 2015; Nouraee and Mowla, 2015) and neurological diseases (Leidinger et al., 2013; Christensen and Schratt, 2009). Other studies have suggested that miRNAs can also participate in intercellular communication (Turchinovich et al., 2019; Baranova et al., 2019).

There is a large number of reports consistently demonstrating the role of miRNAs in viral infections. One of the research directions is connected with miRNAs which can target viral RNAs. Since RNA of single-stranded RNA virus (ssRNA virus) is not structurally distinguishable from host mRNA there are no barriers for miRNA to bind it. In contrast to conventional binding to the 3’-UTR of the target mRNA, host miRNAs often bind to the coding region or to the 5’-UTR of the viral RNA (Bruscella et al., 2017). Besides translational repression such interactions can also enhance viral replication or purposefully alter the amount of free miRNA in a cell (Trobaugh and Klimstra, 2017). For example, miR-122 can bind to the 5’-UTR of the hepatitis C virus (HCV) RNA which increases RNA stability and viral replication since it becomes protected from a host exonuclease activity (Shimakami et al., 2012). Another report contains evidences that miR-17 binding sites on the RNA of bovine viral diarrhea virus (BVDV) are aimed to decrease level of free miR-17 in the cell therefore mediating expression of miRNA targets (Scheel et al., 2016).

Other research groups focused on miRNAs altering their expression in response to the viral infection. Thus, Liu et al. showed that proteins of avian influenza virus H5N1 cause the upregulation of miR-200c-3p in the lungs (Liu et al., 2017). This miRNA targets the 3’-UTR of *ACE2* mRNA therefore decreasing its expression. On the other hand, it was shown that the decrease in *ACE2* expression is critical in ARDS pathogenesis (Imai et al., 2005). Thus, H5N1 virus promotes miRNA-mediated *ACE2* silencing to induce ARDS. Recent reports suggest several other host miRNAs which can potentially regulate *ACE2* and *TMPRSS2* expression which can be also important during SARS-CoV-2 infection due to crucial role of these enzymes for virus cell entry (Nersisyan et al., 2020). Another example was highlighted by Choi et al. who studied miRNAs altering their expression during influenza A virus infection (Choi et al., 2014).

It was shown that several miRNAs which play an important role in cellular processes including immune response and cell death exhibited significant expression differences in infected mice. In the same work authors show that treatment with the respective anti-miRNAs demonstrates an effective therapeutic action.

In recent work, Fulzele with co-authors found hundreds of miRNAs which can potentially bind to the SARS-CoV-2 RNA as well as the RNA of the highly similar SARS-CoV coronavirus (Fulzele et al., 2020). However, this large miRNA list should be narrowed to find high-confidence interactions which should be further experimentally validated. In this work we hypothesized that there can be miRNA-mediated virus-host interplay mechanisms common for several human coronaviruses. For that we used bioinformatic tools to predict miRNA binding sites within human coronavirus RNAs including ones inducing severe acute respiratory syndrome (SARS-CoV-2, SARS-CoV and MERS-CoV) as well as other human coronaviruses causing common cold, namely, HCoV-OC43, HCoV-NL63, HCoV-HKU1 and HCoV-229E. To find and explore more complex regulatory mechanisms we also analyzed miRNome of mouse lungs during SARS-CoV infection to select miRNAs whose expression was significantly altered upon viral infection.

## RESULTS

### Human coronavirus RNAs have a large number of common host miRNA binding sites

To identify human miRNAs that may bind to RNAs of human coronaviruses we used two classical miRNA target prediction tools: miRDB and TargetScan. TargetScan results can be ranked with different seed-region binding types while miRDB results can be ranked with so-called “target score” associated with the probability of successful binding. Interestingly, for each of viruses TargetScan predicted 2-3 times higher number of miRNAs, while 80-85% miRNAs predicted by miRDB were predicted by TargetScan too (summary on number of miRNAs predicted for each of viral genomes is presented in Supplementary File 1).

We constructed the list of 19 miRNAs potentially targeting multiple viral RNAs by selecting miRNAs with miRDB target scores greater than 75 in all analyzed viruses (Supplementary File 1). For the further analysis we selected only high confidence miRNAs according to miRBase, namely, hsa-miR-21-3p, hsa-miR-195-5p, hsa-miR-16-5p, hsa-miR-3065-5p, hsa-miR-424-5p and hsa-miR-421. Target scores of selected miRNAs as well as corresponding hierarchical clustering of viruses are illustrated in Figure 1A. As it can be seen, such clustering grouped together SARS-CoV and SARS-CoV-2 as well as HCoV-229E and HCoV-NL63 which can be also observed when clustering is performed based on the viral genomic sequences similarity (Zhou et al., 2020b). In order to assess which of these miRNAs could demonstrate activity in human lungs we analyzed miRNA sequencing (miRNA-seq) data from The Cancer Genome Atlas Lung Adenocarcinoma (TCGA-LUAD) project. Two of the enlisted miRNAs demonstrated relatively high expression, see Figure 1B. Specifically, hsa-miR-21-3p and hsa-miR-16-5p corresponded to top-5% of the most highly expressed miRNAs according to their mean expression level taken across all samples.

**Figure 1.**
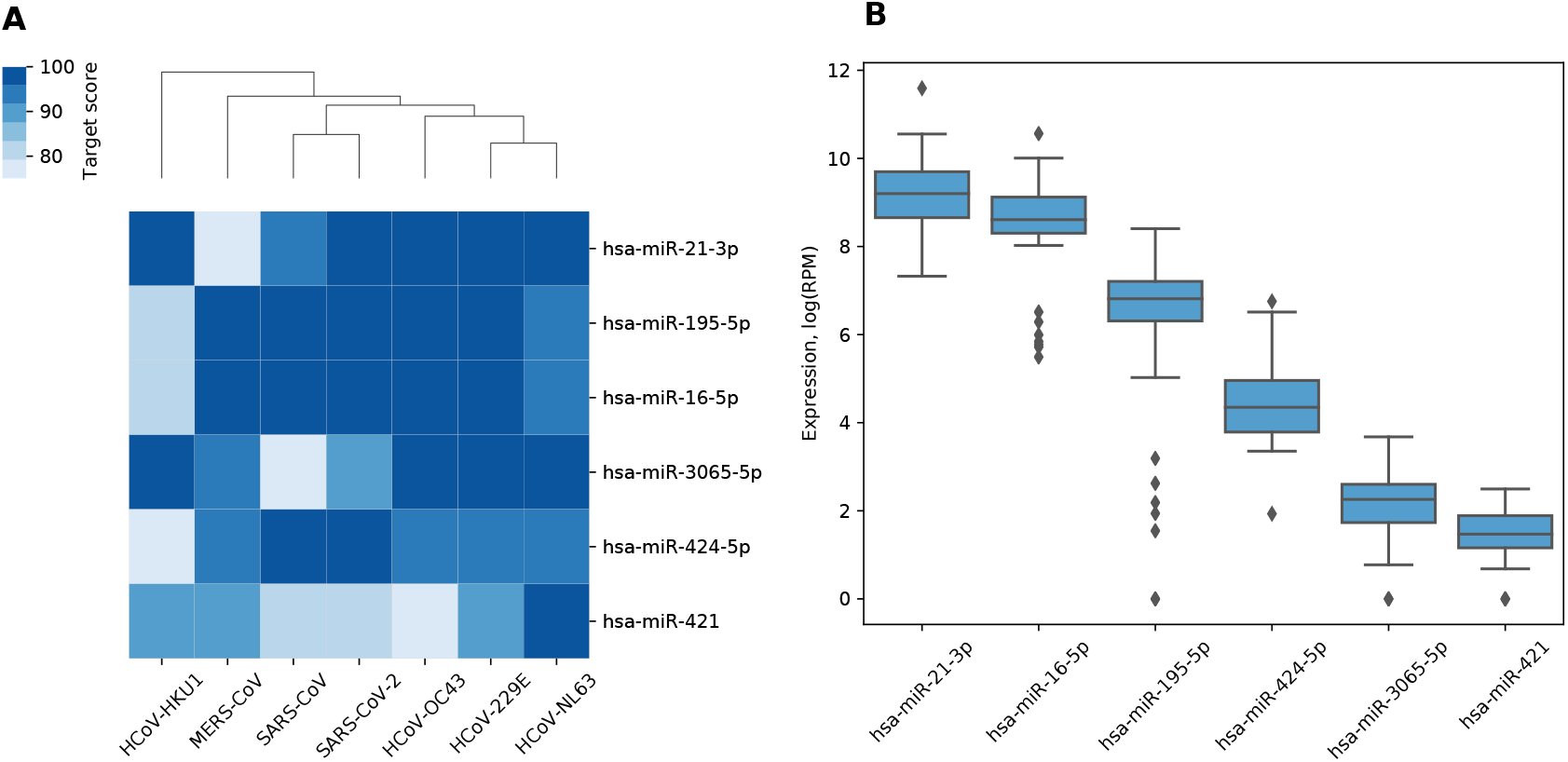
miRNAs with the highest target scores. (A) Hierarchical clustering of coronaviruses based on the miRDB target scores. Rows are sorted according to the mean target score. (B) Expression distribution of miRNAs in human lungs.

### The miR-21 and its target genes exhibit significant expression alteration in mouse lungs during SARS-CoV infection

To further explore a potential interplay between host miRNAs and coronaviruses we hypothesized that some of miRNAs predicted to bind viral RNAs can have altered expression during the infection. In order to test this hypothesis we analyzed two publicly available miRNA-seq datasets of mouse lungs during SARS-CoV infection. The first dataset (GSE36971) included data derived from four mouse strains infected by SARS-CoV and four corresponding control mice. The second dataset (GSE90624) comprised three infected and four control mice.

Differential expression analysis revealed 19 miRNAs in the first dataset and 21 in the second dataset whose expression change during infection was statistically significant (Supplementary File 1). Six miRNAs were differentially expressed in both datasets, five of them had matched fold change signs, namely, were overexpressed in infected mice, see Figure 2. This was a statistically significant overlap since an estimate of the probability for 19- and 21-element random miRNA sets having five or more common elements was equal to 4.07 × 10 ^−7^ (hypergeometric test). Surprisingly, miR-21a-3p which we previously identified as a potential regulator of all analyzed coronavirus genomes with one of the highest scores exhibited 8.3-fold increase (adjusted *p* = 5.72 × 10^−35^) and 11-fold increase (adjusted *p* = 5.77 × 10^−11^) during SARS-CoV infection in the first and the second datasets, respectively.

**Figure 2.**
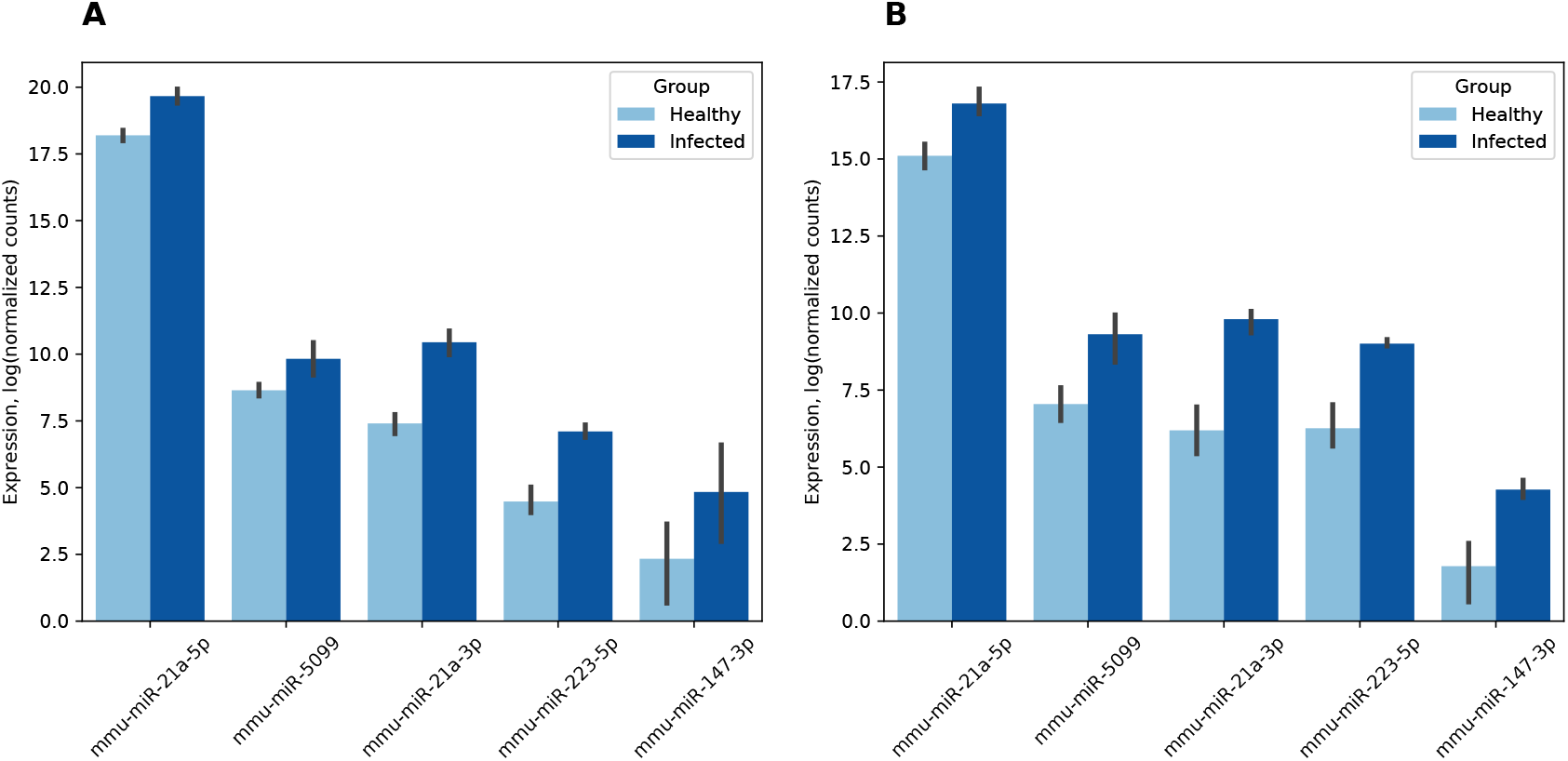
Differentially expressed mouse lung miRNAs. (A) GSE36971. (B) GSE90624. Note that counts were normalized using DESeq2 independently for each dataset. Thus, presented values should not be directly compared across (A) and (B).

Expression of mmu-miR-21a-5p (the opposite “guide” strand of the same hairpin) was also increased in the infected group (2.8- and 3.2-folds, respectively). The latter had a particular importance since mmu-miR-21a-5p was very highly expressed in mice during both experiments. Namely, according to its mean expression across all samples it was 4th and 38th out of 2302 in the first and the second datasets, respectively. Thus, significant expression change of this miRNA can dramatically affect expression of its target genes.

In order to capture aberrant expression of miRNA target genes during infection we analyzed RNA sequencing (RNA-seq) data of eight SARS-CoV infected mice strains published by the same group of authors as in the first miRNA-seq dataset (GSE52405). Two strategies were utilized to generate a list of miRNA targets. Namely, we used the target prediction tools described in the previous section as well as the literature-curated miRTarBase database.

First, we took genes predicted both by miRDB and TargetScan with miRDB target score greater than 75. Additionally, we thresholded this list using top 10% predictions based on TargetScan’s cumulative weighted context++ score. A significant fraction of mmu-miR-21a-5p target genes were down-regulated during the infection. Namely, 6 out of 24 considered genes demonstrated significant decrease in expression (hypergeometric test *p* = 7.6 × 10^−3^). For four other miRNAs there was no statistical significance on the number of down-regulated target genes. Situation was quite different for interactions enlisted in miRTarBase. Thus, 2 out of 2 mmu-miR-21a-3p target genes (*Snca* and *Reck*) were down-regulated (hypergeometric test p = 5.7 × 10^−3^) while only 6 out of 37 mmu-miR-21a-5p target genes exhibited expression decrease (hypergeometric test *p* = 0.057). As in the previous case, no significant number of down-regulated target genes was observed for other miRNAs.

### Viral binding sites of miR-21-3p are conserved across different coronaviruses and their strains

Since miR-21-3p was overexpressed in mouse lung during SARS-CoV infection and was predicted to bind several coronavirus RNAs it is important to analyze the putative binding sites. As summarized in Table 1, all viruses had tens of miR-21-3p binding regions while the peak was observed for HCoV-NL63. To go deeper and analyze mutual arrangement of these sites we performed multiple sequence alignment on seven analyzed genomes and mapped the predicted miR-21-3p binding positions from individual genomes to the obtained alignment. Then, for each binding site mapped to the alignment we calculated the number of viruses sharing that particular miRNA binding site.

**Table 1.** Number of miR-21-3p binding sites on coronavirus RNAs. Seed region binding types are named according to TargetScan. The last column indicates the number of binding sites having not a single mutation in analyzed viral strains.

|  | 6mer | 7mer-1a | 7mer-m8 | 8mer-1a | Total | Without mutations |
| --- | --- | --- | --- | --- | --- | --- |
| SARS-CoV-2 | 13 | 3 | 17 | 2 | 35 | 27 |
| SARS-CoV | 9 | 4 | 8 | 3 | 24 | 17 |
| MERS-CoV | 8 | 1 | 7 | 1 | 17 | 11 |
| HCoV-OC43 | 9 | 3 | 13 | 7 | 32 | 20 |
| HCoV-NL63 | 21 | 7 | 24 | 12 | 64 | 55 |
| HCoV-HKU1 | 12 | 7 | 14 | 8 | 41 | 34 |
| HCoV-229E | 10 | 4 | 18 | 11 | 43 | 36 |

In total, there were 34 positions common for two or more viruses. Interestingly, 31 of them corresponded to nonstructural proteins located in the polyprotein 1ab coding region. Two other positions were located in spike protein, both were shared by HCoV-HKU1 and HCoV-OC43. Besides, one position was located in the nucleocapsid protein of HCoV-229E and HCoV-NL63. One position on the alignment was specific to six out of seven considered coronaviruses, namely, a consensus was obtained for miR-21-3p binding sites on polyprotein 1a of all viruses except MERS-CoV, see Figure 3. One other position corresponded to five viruses (SARS-CoV-2, SARS-CoV, HCoV-HKU1, HCoV-229E, HCoV-NL63), while two positions were shared by four viruses (SARS-CoV-2, SARS-CoV, HCoV-229E, HCoV-NL63) and six positions were shared by three viruses. Full information is listed in Supplementary File 1.

**Figure 3.**
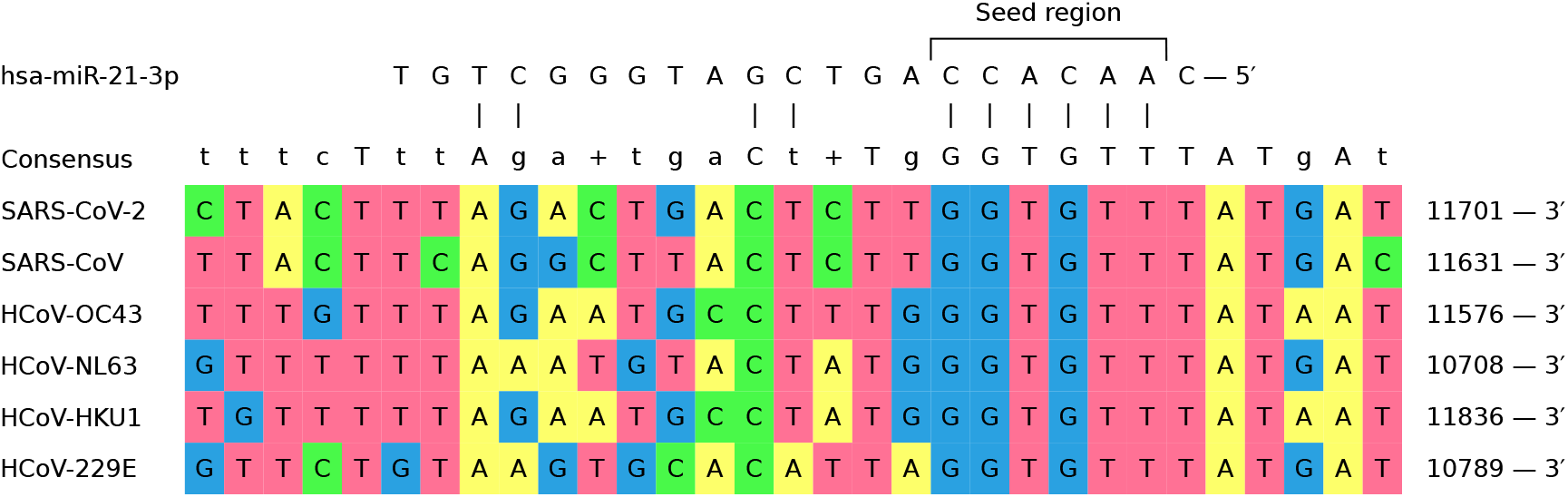
The shared binding site of miR-21-3p on six out of seven human coronaviruses.

To group coronaviruses based on the probability of sharing common binding positions of miR-21-3p we calculated the number of matching positions of binding sites in the multiple alignment for each pair of viruses, see Figure 4A. This data was normalized by 34 (total number of binding positions in the alignment shared by two or more viruses) and used as a distance matrix for hierarchical clustering, see Figure 4B. Interestingly, such clustering fully agreed with the clustering of viruses based on their genomic sequence similarity (Zhou et al., 2020b).

**Figure 4.**
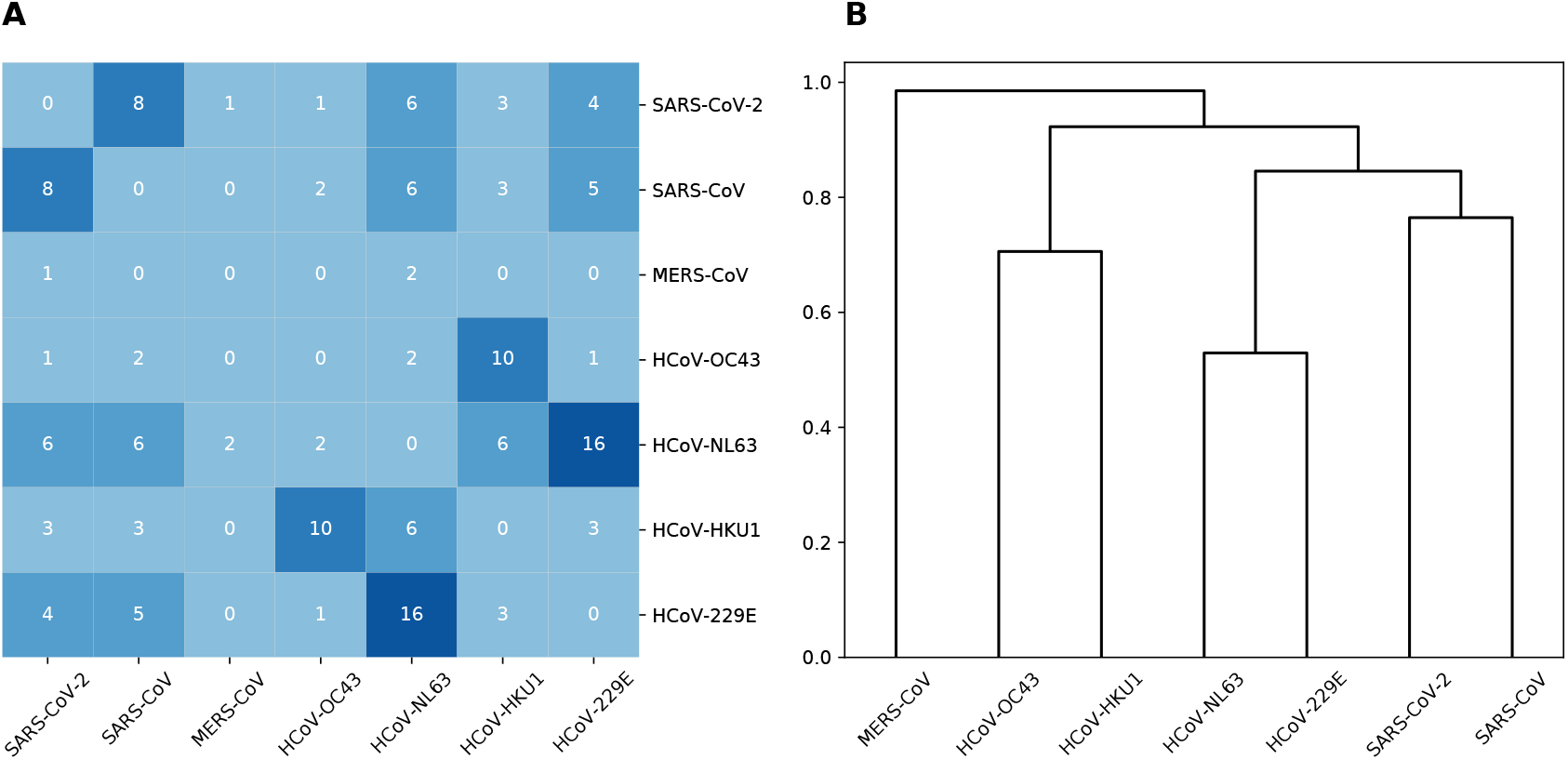
Shared miR-21-3p binding sites. (A) Number of shared binding sites for each pair of human coronaviruses. (B) Hierarchical clustering of coronaviruses based on the number of shared miR-21-3p binding sites.

Finally, to assess conservativity of miR-21-3p binding regions across viral strains we performed multiple sequence alignment of available viral genomes independently for each of human coronaviruses. The results demonstrated high conservativity of these regions: 63% to 86% of binding sites within coronavirus RNAs had not a single mutation (see Table 1) while mean of average mutation rates (i.e. number of mutations normalized by region length and number of strains) across all viruses was equal to 0.5%. Even more interesting results were obtained for the already mentioned binding sites common for several viruses. Namely, for 24 out of 34 regions (71%) there were no mutations in each of viruses, including the one shared by six coronaviruses. For mutated regions average mutation rate was limited by 5%, mean was equal to 0.9%. The exact values of average mutation rates are available in Supplementary File 1.

## DISCUSSION

In this work we found that miRNA miR-21-3p has a potential to bind multiple human coronavirus RNAs. Moreover, aside from virus specific binding sites we identified genomic positions which can serve as conserved miR-21-3p targets for several RNAs. As one can expect, viruses with high genomic similarity such as SARS-CoV-2 / SARS-CoV or HCoV-229E / HCoV-NL63 had higher number of shared binding sites.

On the other hand, we found that expression of miR-21-3p in lungs exhibits an 8-fold increase upon SARS-CoV infection. Interestingly, miR-21-5p (the “guide” strand of the same pre-miRNA hairpin) demonstrated only 3-folds increase in expression. To explain this phenomena of non-symmetrical expression increase we hypothesize that binding to the viral genome saves star miRNA miR-21-3p from degradation after unsuccessful attempt of AGO2 loading. A similar mechanism was already mentioned in several works. Namely, Janas with co-authors demonstrated that Ago-free miRNAs can escape degradation by forming Ago-free miRNA-mRNA duplex (Janas et al., 2012) thereby confirming previously proposed hypotheses (Song et al., 2003; Chatterjee and Großhans, 2009). A similar concept was named as target RNA directed miRNA degradation (TDMD) which consists of the fact that highly complementary target RNA can trigger miRNA degradation by a mechanism involving nucleotide addition and exonucleolytic degradation (la Mata et al., 2015). This concept was experimentally validated as an example of miR-574-5p and miR-574-3p study in gastric cancer (Zhang et al., 2019). Thus, non-proportional up-regulation of miR-21 arms can be indirect evidence that miR-21-3p directly targets the viral RNA or that the miR-21-5p is being actively degraded during target mRNA binding.

Several hypotheses can be put forward to explain biological motivation of such interplay. At first sight, one can think about host miRNA-mediated immune response to the viral infection. For example, translation of human T cell leukemia virus type I (HTLV-1) is inhibited by miR-28-3p activity (Bai and Nicot, 2015). However, our results suggest that miR-21-3p binding sites are actually conserved across human coronaviruses. Thus, viruses can purposefully accumulate host miRNA binding sites to slow down their own replication rate in order to evade fast detection and elimination by the immune system. Such behaviour was reported e.g. in the case of eastern equine encephalitis virus (EEEV) (Trobaugh et al., 2014). Authors reported that haematopoietic-cell-specific miRNA miR-142-3p directly binds viral RNA which limits the replication of virus thereby suppressing innate immunity. The latter was shown to be crucial in the virus infection pathogenesis.

A functional activity of miR-21-3p was already mentioned in the context of viral infections. Thus, it was shown that miR-21-3p regulates the replication of influenza A virus (IAV) (Xia et al., 2018). Namely, hsa-miR-21-3p targeting 3’-UTR of *HDAC8* gene was shown to be down-regulated during IAV infection of human alveolar epithelial cell line A549 using both miRNA microarray and quantitative PCR analysis. Consecutive increase in the *HDAC8* expression was shown to promote viral replication.

Indeed, overexpression of miRNA can lead to down-regulation of its target genes. Multiple reports indicate a role of miR-21-3p in such pathways as cell proliferation, cell cycle phases, and the DNA metabolic process. Gao et al. demonstrated that the miR-21-3p can promote cell proliferation and decrease apoptosis in cancer stem cells (CSCs) by regulating *TRAF4*, which functions are related to cell proliferation and apoptosis (Gao et al., 2019). In another work Liu et al. demonstrated that the downregulation of miR-21-3p can inhibit the apoptosis caused by lipopolysaccharide (LPS) (Liu et al., 2019). The apoptosis inhibition was shown to happen due to the targeting of the 3’-UTR of *RGS4* mRNA by miR-21-3p. There are also prognostic signatures based on the level of miR-21-3p expression. Namely, hsa-miR-21-3p together with *L1CAM* gene form a reliable pair for overall and disease free survival in ovarian, endometrial, breast, renal cell carcinoma and pancreatic ductal adenocarcinoma (Doberstein et al., 2014). Finally, miR-21-3p was shown to control sepsis-associated cardiac dysfunction by targeting *SORBS2* gene (Wang et al., 2016).

## MATERIALS AND METHODS

### Prediction of miRNA binding sites

To find miRNAs which can bind to the viral RNAs we used miRDB v6.0 (Chen and Wang, 2020) and TargetScan v7.2 (Agarwal et al., 2015). Viral genomes and their annotations were downloaded from the NCBI Virus (Hatcher et al., 2017) under the following accession numbers:

- NC_045512.2 (SARS-CoV-2);
- NC_004718.3 (SARS-CoV);
- NC_019843.3 (MERS-CoV);
- NC_006213.1 (HCoV-OC43);
- NC_005831.2 (HCoV-NL63);
- NC_006577.2 (HCoV-HKU1);
- NC_002645.1 (HCoV-229E).

For the analysis of miRNA-mRNA interactions we also used miRTarBase v8 (Huang et al., 2020).

### RNA sequencing data and differential expression analysis

The miRNA-seq data from the TCGA-LUAD project (Collisson et al., 2014) was used to quantify miRNA expression in the human lungs. Specifically, the data was obtained from GDC Data Portal (https://portal.gdc.cancer.gov/) and included miRNA expression table whose columns correspond to 46 normal lung tissues and rows are associated with miRNAs. We used log_2_-transformed Reads Per Million mapped reads (RPM) as a miRNA expression unit.

Two miRNA-seq datasets, GSE36971 (Peng et al., 2011) and GSE90624 (Morales et al., 2017), were used to analyze miRNome of SARS-CoV infected mouse lungs. Raw fastq files were downloaded from Sequence Read Archive (Leinonen et al., 2011). Adapters were trimmed via Cutadapt 2.10 (Martin, 2011), miRNA expression was quantified by miRDeep2 (Friedländer et al., 2012) using GRCm38.p6 mouse genome (release M25) from GENCODE (Frankish et al., 2019) and miRBase 22.1 (Kozomara et al., 2019). Gene expression profile of SARS-CoV infected mouse lungs was downloaded in form of count matrix from Gene Expression Omnibus (GEO) (Barrett et al., 2013) under GSE52405 accession number (Josset et al., 2014). Differential expression analysis was performed with DESeq2 (Love et al., 2014). The results were filtered using 0.05 threshold on adjusted p-value and 1.5 on fold change (linear scale).

### Sequence alignment

Multiple Sequence Alignment (MSA) of viral genomic sequences was done using Kalign 2.04 (Lassmann et al., 2009). Two MSA series were performed. In the first one we aligned seven human coronavirus genomes. In the second one different coronavirus strains were aligned for each of analyzed viruses. All genomes available on the NCBI Virus were used for SARS-CoV, MERS-CoV, HCoV-OC43, HCoV-NL63, HCoV-HKU1 and HCoV-229E (263, 253, 139, 58, 39 and 28 genomes, respectively). For SARS-CoV-2 thousand genomes were randomly chosen preserving the percentage of samples from each country. For each virus we established the mapping between alignment and genomic coordinates. With the use of this mapping, miRNA seed region binding positions within viral RNAs were placed on the alignment.

## Data and code availability

All code was written in Python 3 programming language with extensive use of NumPy (Van Der Walt et al., 2011) and Pandas (McKinney, 2010) modules. Statistical analysis was performed using the SciPy stats (Virtanen et al., 2020), plots were constructed using the Seaborn and Matplotlib (Hunter, 2007). MAS was visualized using Unipro UGENE (Okonechnikov et al., 2012). All used data and source codes are available on GitHub (https://github.com/s-a-nersisyan/host-miRNAs-vs-coronaviruses).

## ACKNOWLEDGMENTS

The authors thank Dr. Maxim Shkurnikov for valuable comments and discussions.

